# Matching a snail’s pace: Successful use of environmental DNA techniques to detect early stages of invasion by the destructive New Zealand mud snail

**DOI:** 10.1101/2020.08.12.248948

**Authors:** James D. Woodell, Maurine Neiman, Edward P. Levri

## Abstract

Early detection of invasive species allows for a more rapid and effective response. Restoration of the native ecosystem after an invasive population has established is expensive and difficult but more likely to succeed when invasions are detected early in the invasion process. Containment efforts to prevent the spread of known invasions also benefit from earlier knowledge of invaded sites. Environmental DNA (eDNA) techniques have emerged as a tool that can identify invasive species at a distinctly earlier time point than traditional methods of detection. Due to expected range expansion in eastern North America, we focus on the destructive New Zealand Mud Snail *Potamopyrgus antipodarum* (NZMS) invasion. We collected water samples from eight sites that prior evidence indicated were not yet invaded by the NZMS. After filtering these samples to collect eDNA, we used a species-specific probe with qPCR to identify NZMS eDNA. We found evidence for NZMS invasion at five of the eight sites, with later physical confirmation of mud snails at one of these sites. This study is the first example of successful detection of a previously unidentified invasive population of NZMS, setting the stage for further monitoring of at-risk sites to detect and control new invasions of this destructive snail. This study also shows potential opportunities for invasion monitoring offered by using low-cost efforts and methods that are adaptable for citizen science.

## INTRODUCTION

Halting initial introductions has been identified as the surest and most cost-effective method of invasion mitigation (Finoff *et al*. 2007, Keller *et al*. 2007, NISC 2016). Challenges to the effective implementation of invasion prevention are posed by absence of sufficient policy or because the opportunity to prevent invasion has already passed (Simberloff 2014). The next priority should then be to eliminate the invader completely before greater damage and further spread can occur (Simberloff *et al*. 2013). While eradication is increasingly possible (Simberloff 2014), restoration of invaded ecosystems is time consuming, expensive, and in many cases unsuccessful (Myers *et al*. 2000, Rejmánek and Pitcairn 2002). Accordingly, current invasive species response efforts often focus on preventing spread rather than eradication (Leung *et al*. 2002). Establishing cost- and time-effective methods to first identify and then stop or slow the spread of early invasions is a critical means of preventing substantial future biological and economic damage (Lodge *et al*. 2006, McGeoch *et al*. 2015).

Perhaps the most important element of a rapid response strategy is identification of a new invasion (Simberloff 2014, McGeoch *et al*. 2015). Traditional methods of surveying ecosystems for invasion requires the species to be physically located (Lawson Handley 2015), meaning labor-intensive field sampling (and a little luck). By the time a site is acknowledged as invaded, the invader is often well established and has likely spread to other sites (Simberloff 2014). This outcome is particularly likely when active searches are not regularly performed and discovery of the invasion occurs via chance encounters. This lag time between actual invasion and realization that an invasion has occurred introduces a critical period during which an unaware human population may help spread the invasive species because containment measures cannot be initiated until we are aware of the invasion. Early detection is also important because full remediation is more likely to succeed if the invasion is caught in its early stages (Rejmanek and Pitcairn 2002, Anderson 2005, Simberloff 2014, U.S. Department of the Interior 2016). Decreasing this lag time associated with invasive species containment by focusing on early detection of invasions should thus be a key priority for invasive species management by conservation programs.

Traditional methods of detecting new invasive populations often fail to catch invasions during the early stages of invasion when the ability to contain or eradicate these populations is maximized (U.S. Department of the Interior 2016). Environmental DNA (eDNA)-based approaches have emerged as a promising means to monitor ecosystems for the introduction of invasive species during the establishment process (Jerde *et al*. 2011, Taberlet *et al*. 2012, Comtet *et al*. 2015, Lawson Handley 2015, Thomsen and Willerslev 2015, Brown *et al*. 2016, Ricciardi *et al*. 2017, Jerde 2019, Sepulveda *et al*. 2020).

### Early Detection Using Environmental DNA

Environmental DNA (eDNA), defined as genetic material sampled from the environment rather than directly from organisms, is rapidly emerging as a powerful means of surveying natural biological communities (Jerde *et al*. 2011, Taberlet *et al*. 2012, Comtet *et al*. 2015, Lawson Handley 2015, Thomsen and Willerslev 2015, Brown *et al*. 2016, Ricciardi *et al*. 2017). For conservation purposes, eDNA can detect the presence of species where traditional surveying did not (Wilcox *et al*. 2016, Smith and Goldberg 2019), including early identification of previously unreported populations (Brown *et al*. 2016).

Sources of eDNA include soil, water, sediment, or any medium that can harbor DNA from a living or recently dead organism. As an organism moves within an environment, it sloughs off skin cells or leaves behind wastes that contain its DNA. After taking samples of the environment, the sample is processed to isolate and amplify the DNA contained within the nonorganic components. This DNA is then used to describe the metagenomics of the community and/or identify species within that community (Comtet *et al*. 2015).

While initially developed for surveying microbial communities, eDNA is now used for various ecological surveys, including macro-organisms. There are now multiple species-specific assays available for use, including assays for numerous amphibians (Beauclerc *et al*. 2018), fish (Thomsen *et al*. 2012), plants (Scriver *et al*. 2015), and mammals (Andersen *et al*. 2012), and more assays are continually being developed (Thomsen and Willerslev 2015, see Washington State University’s eDNA toolbox: https://labs.wsu.edu/edna/edna-assays/). Environmental DNA techniques have also proven capable of detecting invasive species where traditional surveys had not (Jerde *et al*. 2011).

For invasive species conservation, the application of eDNA-based approaches allows a more efficient means of surveying potentially invaded sites for invasive species. In particular, these surveys no longer require physical location of an organism. For example, eDNA has been used to detect invasive or endangered fish, frogs, and crustaceans in aquatic ecosystems without the need to physically locate individuals at those sites (see Jerde *et al*. 2011, Dejean *et al*. 2012, Takahara *et al*. 2013). Instead, samples of the environment can be acquired and processed at low cost and used in conjunction with a species-specific probe in order to identify the presence or absence of an invader in ecosystems of concern. Environmental DNA can therefore detect invasive species at a much earlier time point, allowing substantially more rapid management and remediation responses. Because eDNA rapidly degrades in the external environment, identifying DNA from an organism means that organism has recently been present in that ecosystem. Diffusion of eDNA, particularly in aquatic ecosystems, may also allow for detection of invaders beyond the site of deposition (*e*.*g*., downstream from the invasion site; Dejean *et al*. 2011, Deiner and Altermatt 2014). Calibrating quantitative PCR (qPCR) probe fluorescence with known population densities and river discharge or water volume also makes it possible to estimate invasive population densities in the aquatic environments (Goldberg *et al*. 2013). Here, we describe the first successful application of eDNA to detect a previously unknown population of the destructive invasive New Zealand snail *Potamopyrgus antipodarum*. We also provide an example of the efficacy of eDNA monitoring as compared to traditional surveys for identifying invasive species in at-risk areas.

### New Zealand Mud Snail Invasion

*Potamopyrgus antipodarum*, commonly called the New Zealand Mud Snail (NZMS) in invaded regions, is native to freshwater lakes, streams, and (rarely) estuarine environments in New Zealand. NZMS was first observed in the River Thames in the 1850s (Smith 1889). In 1892, NZMS was recorded in Tasmania, and was found in mainland Australia in 1895 (Alonso and Castro-Díez 2008). Soon after the turn of the 20th century, NZMS was discovered in brackish waters in northern Europe (Städler *et al*. 2005). NZMS became a prominent invader later in the 20th century, establishing inland in France and Switzerland in the 1970s and then expanding across the rest of central Europe (Städler *et al*. 2005). Invasion of North America by NZMS was discovered in 1987 in the Snake River in Idaho (Bowler 1991). The New Zealand Mud Snail has subsequently expanded along rivers and lakes of the western US, including sites in Colorado (McKenzie *et al*. 2013), Utah (Vinson 2004), Wyoming (Kerans *et al*. 2005), Washington (Davidson *et al*. 2008), and California and Oregon (Dybdahl and Drown 2011). The western invasive NZMS populations have also expanded into Canada in brackish waters along the Pacific coast, similar to the invasion of brackish waters in Northern Europe (Davidson *et al*. 2008). An additional independent invasion of North America occurred first detected in Lake Ontario and the St. Lawrence River in 1991 (Zaranko *et al*. 1997), likely a secondary invasion from European invasive populations (Dybdahl and Drown 2011, Donne *et al*. 2020). New Zealand mud snail has since spread across the Great Lakes (Levri and Jacoby 2008, Levri *et al*. 2007, 2008, 2012) and other watersheds in the Eastern US in New York (R. Hood, pers comm.), Pennsylvania (R. Morgan, pers comm., Levri *et al*. 2020), and Maryland (J. Kilian, pers. comm.). Additional documented invasions around the world include the Black Sea (Son 2008), Italy (Gaino *et al*. 2008), Japan (Ogata *et al*. 2010), South America (Collado and Fuentealba 2020), Spain and Portugal (Alonso *et al*. 2019), and Turkey (Odabaşi *et al*. 2019). NZMS native to New Zealand are either sexual or asexual, but invasive populations seem to be invariably asexual (Alonso and Castro-Díez 2012). While asexual NZMS harbor a great deal of genetic diversity (Jokela *et al*. 2003), only a handful of NZMS clones have been successful invaders (Städler *et al*. 2005, Alonso and Castro-Díez 2012, Donne *et al*. 2020).

Multiple studies have provided important insights into the current and potential consequences of NZMS invasion. First, invasive NZMS populations can grow to extremely high densities, exceeding 500,000 individuals•m^-2^ (Hall *et al*. 2006, Tatara *et al*. 2012). This physical density can translate into the loss, via competitive exclusion, of other species that colonize or dwell along the substrate (Alonso and Castro-Díez, 2012). For example, experimental studies demonstrate a negative effect of NZMS on macroinvertebrate colonization where NZMS populations are relatively high (Kerans *et al*. 2005). Other experiments have demonstrated that rainbow trout (*Oncorhynchus mykiss*) fed exclusively NZMS lose weight because the fish are not able to digest the snails (Vinson and Baker 2008), and NZMS are a poorer food source than other gastropods for tench (Butkus and Višinskienė 2020). The implications are that NZMS invasions have serious potential consequences that could affect multiple trophic levels in invaded ecosystems, especially in light of the observation that native fish are increasingly consuming NZMS (Vinson and Baker 2008).

The New Zealand Mud Snail has had demonstrably negative effects on ecosystems where they have successfully established. Hall *et al*. (2003) showed that invasive NZMS dominated nitrogen and carbon fluxes in the invaded Polecat Creek in Wyoming, finding that NZMS consumed 75% of gross primary productivity, represented two-thirds of ammonium demand, and constituted 97% of invertebrate biomass. Krist and Charles (2012) discovered that invasive NZMS also seem to outcompete native grazers, perhaps via direct competition for food. This study also revealed that competition imposed by NZMS altered native diatom community assemblages. Diatom communities can be used as ecological indicators for environmental conditions, meaning changes in diatom community structures are often indicative of environmental changes (Keck *et al*. 2016, Pandey *et al*. 2018). Moore *et al*. (2012) found that invasive NZMS, which use their radula to scrape algae off substrate, altered algal communities via direct competition with native scraping grazers, reporting an increase in piercing-type grazers in the community from 0 individuals•m^-2^ to an average of 1500 individuals•m^-2^. This shift from scraping to piercing-type grazers is associated with depleted stable nitrogen isotopes in native invertebrates (Moore *et al*. 2012). Community phase shifts are indicative that NZMS are dramatically altering the ecosystems they invade. As these alterations continue, predicting outcomes, mitigating damages, and restoring the native environment and community will become increasingly difficult.

While the range of the western NZMS invasion is well characterized, the full extent of the NZMS invasion in the eastern US is less defined. NZMS was discovered in Centre County, Pennsylvania in 2013 at Spring Creek, but was well established when discovered and might have persisted undetected for years (Levri *et al*. 2020). Spring Creek is a popular fishing location, raising suspicions that these NZMS were transported via recreational water use. Data pointing in this direction include the genetic background of the Spring Creek population compared to other invasive populations in the US. Mitochondrial data suggest two primary invasive clones in the US: US1 in the western US and US2 in the Laurentian Great Lakes (Dybdahl and Drown 2011). These are likely separate invasions (Donne *et al*. 2020): the US1 haplotype matches haplotype 37 (Genbank AY570216, Neiman and Lively 2004) found in Lake Taupo on the North Island of New Zealand, while the US2 haplotype matches haplotype 22 (Genbank AY570201, Neiman and Lively 2004; 2% sequence divergence from the US1 haplotype, Dybdahl and Drown 2011) found in lakes Gunn and Te Anau at the southeastern tip of New Zealand (Neiman and Lively 2004). The US2 haplotype also matches the invasive European A mitochondrial haplotype (Dybdahl and Drown 2011), indicating a possible secondary invasion originating with the successful invasive population in Europe (also see Donne *et al*. 2020). An additional mitochondrial haplotype was discovered at a single site in lower densities alongside US1, US3 (Genbank HG680431, <0.2% sequence divergence from the US2 mitochondrial haplotype). These US3 snails are not currently considered invasive as they have not been identified beyond a single site and are a minority compared to the greater US1 population at that location (Dybdahl and Drown 2011). The invasive population found in Spring Creek is comprised entirely of a clone with the US1 mitochondrial haplotype (M. Dybdahl, pers. comm.), matching the dominant clone in the western US rather than the US2 haplotype of the geographically nearer Great Lakes populations. Because the western US1 population has existed at least since 1987 (Bowler 1991), the invasion in Pennsylvania is likely a secondary invasion originating via human-mediated transport of individuals from the western US (also see Donne *et al*. 2020).

That the Spring Creek invasive NZMS have the US1 haplotype is concerning given the widespread invasion of this lineage, which currently ranges from California to southern Canada to Colorado (Vinson 2004, Kerans *et al*. 2005, Davidson *et al*. 2008, Dybdahl and Drown 2011, McKenzie *et al*. 2013). The possibility of recreational transport of NZMS poses a threat to the local trout population and raises the potential for NZMS to be accidentally transported to new localities through ballast water or fishing equipment. In particular, it is very plausible that NZMS has already spread to new eastern North American sites where it has established new invasive populations but has remained undetected. Here, we attempted the first application of eDNA-based early detection of NZMS of which we are aware.

## METHODS

Developed at the University of Idaho (Goldberg *et al*. 2013), eDNA and qPCR protocols for NZMS have proven effective at detecting these snails in known invasion sites and at estimating population density in streams with measured discharge. To our knowledge, these methods have not previously been applied to identifying new invasive populations of NZMS. We successfully refined the filtering protocols from Goldberg *et al*. (2013) and then used these updated methods in a stream water survey in central PA in May 2018. We focused on applying eDNA to determine whether NZMS might be found in locations that could be plausibly invaded but where no snails had previously been reported.

### Site Selection

We selected eight sites at risk of recreational aquatic activity-related transport of new colonists and that represented a more significant risk of further human-mediated spread after an invasion occurred (Table 1, Figure 1). The eight selected sites were spread across six different rivers and four counties in central Pennsylvania. The sites all are contained within the Susquehanna River watershed, which ultimately feeds into the Chesapeake Bay as part of the Mid-Atlantic watershed. Due to a lack of stream discharge measurements across these sites and the potential for inaccurate population density estimates (Darling and Blum 2007), we chose to assess only presence/absence of NZMS at these selected sites rather than monitoring eDNA densities. Six of these sites had been the focus of multiple unsuccessful searches for physical evidence of NZMS invasion between 2014 and 2018 (Levri *et al*. 2020); two sites, at Yellow Creek (YC1) and Cedar Run (CR1), had not been searched prior to this study. Time constraints and unusually high water level prevented thorough traditional sweep net-based visual searches for NZMS at these eight sites on the days of sampling.

**Table 1.**
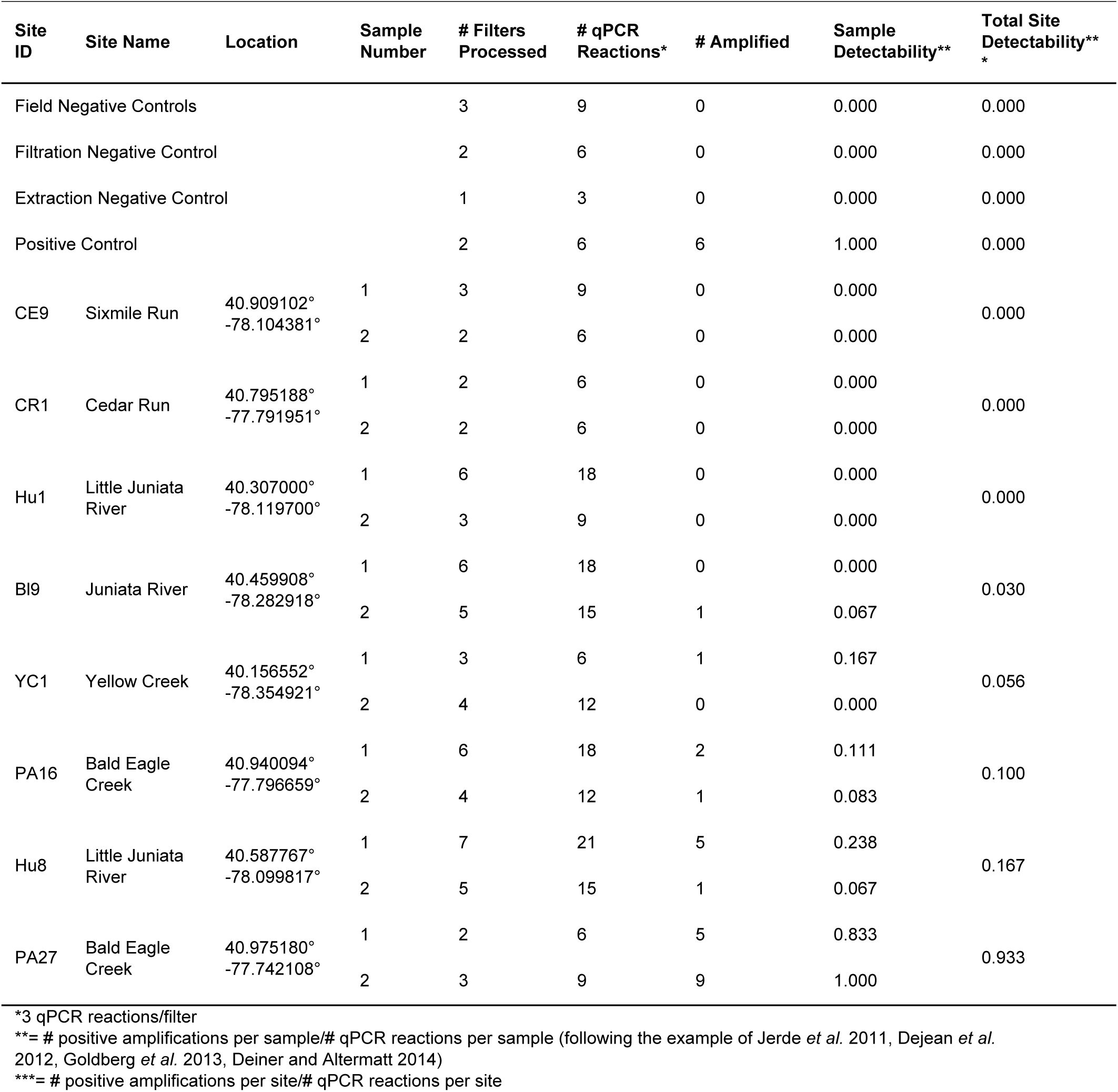
Results of qPCR from sites sampled in May of 2018. All sites sampled were locations where NZMS were not detected previously. Locations of sites can be seen in Figure 1. Positive control consisted of a sample from a lab aquarium containing NZMS. Field negative controls were DI water transported to field sites, transferred to new containers, and filtered as though they were collected samples. Extraction negative control was DI water processed alongside sample filters for DNA extraction. Amplification was considered positive if probe fluorescence reached an exponential phase during the qPCR assay.

| Site ID | Site Name | Location | Sample Number | # Filters Processed | # qPCR Reactions* | # Amplified | Sample Detectability** | Total Site Detectability**<br>* |
| --- | --- | --- | --- | --- | --- | --- | --- | --- |
| Field Negative Controls |  |  |  | 3 | 9 | 0 | 0.000 | 0.000 |
| Filtration Negative Control |  |  |  | 2 | 6 | 0 | 0.000 | 0.000 |
| Extraction Negative Control |  |  |  | 1 | 3 | 0 | 0.000 | 0.000 |
| Positive Control |  |  |  | 2 | 6 | 6 | 1.000 | 0.000 |
| CE9 | Sixmile Run | 40.909102°<br>-78.104381° | 1 | 3 | 9 | 0 | 0.000 | 0.000 |
|  |  |  | 2 | 2 | 6 | 0 | 0.000 |  |
| CR1 | Cedar Run | 40.795188°<br>-77.791951° | 1 | 2 | 6 | 0 | 0.000 | 0.000 |
|  |  |  | 2 | 2 | 6 | 0 | 0.000 |  |
| Hu1 | Little Juniata River | 40.307000°<br>-78.119700° | 1 | 6 | 18 | 0 | 0.000 | 0.000 |
|  |  |  | 2 | 3 | 9 | 0 | 0.000 |  |
| BI9 | Juniata River | 40.459908°<br>-78.282918° | 1 | 6 | 18 | 0 | 0.000 | 0.030 |
|  |  |  | 2 | 5 | 15 | 1 | 0.067 |  |
| YC1 | Yellow Creek | 40.156552°<br>-78.354921° | 1 | 3 | 6 | 1 | 0.167 | 0.056 |
|  |  |  | 2 | 4 | 12 | 0 | 0.000 |  |
| PA16 | Bald Eagle Creek | 40.940094°<br>-77.796659° | 1 | 6 | 18 | 2 | 0.111 | 0.100 |
|  |  |  | 2 | 4 | 12 | 1 | 0.083 |  |
| Hu8 | Little Juniata River | 40.587767°<br>-78.099817° | 1 | 7 | 21 | 5 | 0.238 | 0.167 |
|  |  |  | 2 | 5 | 15 | 1 | 0.067 |  |
| PA27 | Bald Eagle Creek | 40.975180°<br>-77.742108° | 1 | 2 | 6 | 5 | 0.833 | 0.933 |
|  |  |  | 2 | 3 | 9 | 9 | 1.000 |  |

\*3 qPCR reactions/filter
\*\*= # positive amplifications per sample/# qPCR reactions per sample (following the example of Jerde *et al.* 2011, Dejean *et al.* 2012, Goldberg *et al.* 2013, Deiner and Altermatt 2014)
\*\*\*= # positive amplifications per site/# qPCR reactions per site

**Figure 1.**
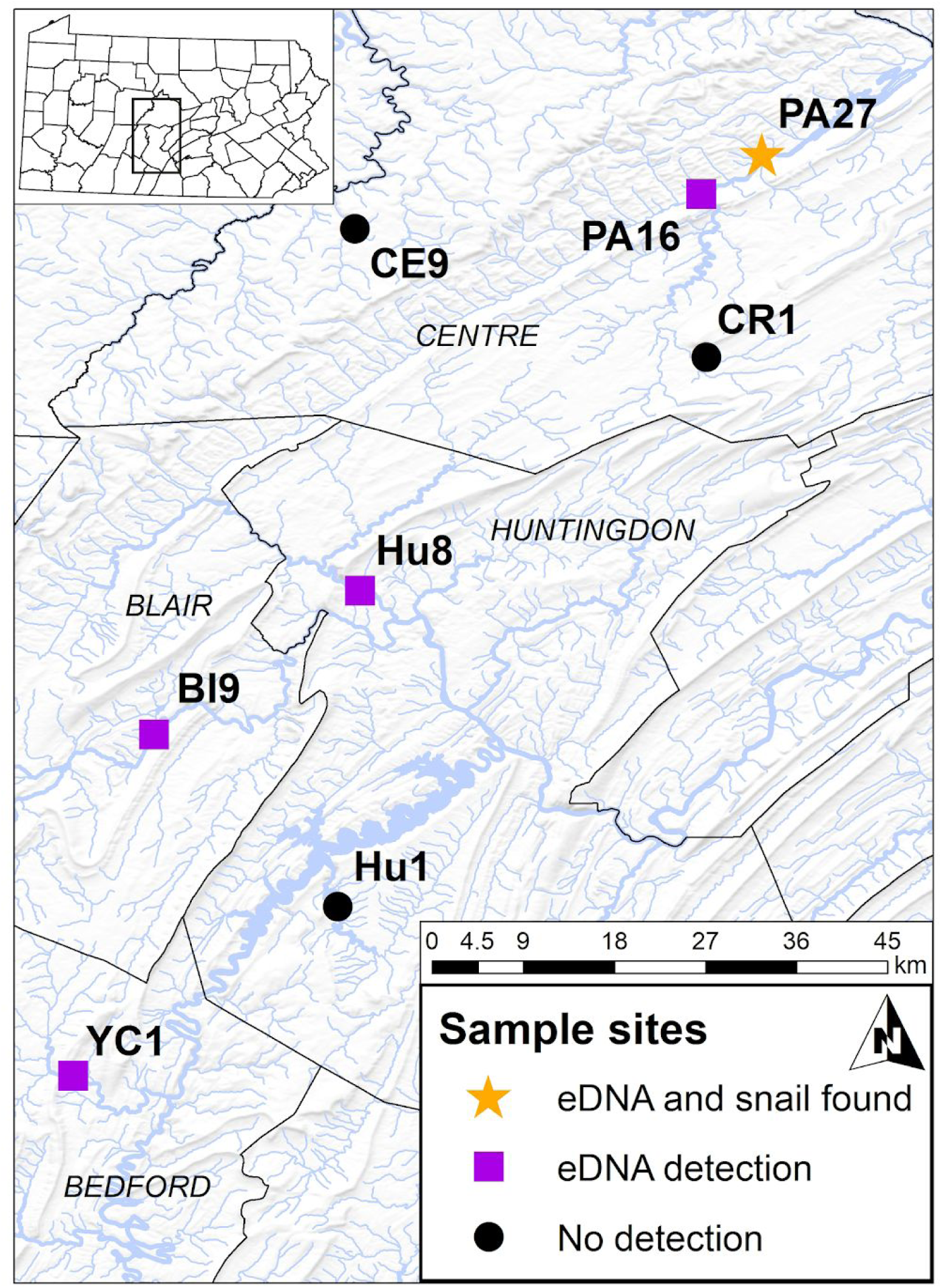
Map of sampling sites in central Pennsylvania. Circles are sites where no NZMS eDNA was found. Squares indicate the presence of eDNA but without confirmation of physical presence. The star marks site PA27 on the Bald Eagle Creek where NZMS eDNA was found and the physical presence of NZMS was later confirmed through hand sampling.

### Field Collections and Filtration

We collected two water samples of 3.8 liters from each of the eight sites by submerging containers approximately 10 cm under the surface of the stream until full. We soaked each container in a 50/50 bleach solution and rinsed the containers thoroughly with deionized water before use. We collected one sample from the bank and one sample from the center of the stream when the waters were relatively shallow and slow moving. In deeper or higher velocity streams, we took two bank samples at two locations moving approximately 10 m upstream for the subsequent sample. For a negative field control, a blank sample consisting of only deionized water was transported to the field along with other containers. This deionized water was then transferred from its original container into a new container after collections were made to check for contamination of samples during transport. We filtered all water samples at the Altoona Campus of the Pennsylvania State University within 24 hours of collection.

We used Nalgene vacuum filter flasks that were sterilized by soaking in a 50/50 bleach and deionized water solution for at least 15 minutes followed by a thorough rinse with deionized water before starting the filtration process for each sample. We also used a 50% bleach solution to sterilize the workspace used for the flasks as well as the forceps that we used for sample processing. As an additional means to prevent contamination, we placed fresh paper towels under the flasks during filtration for each sample. We used 0.45 um mixed-cellulose ester filter discs for filtration. Because of relatively high sediment load in the water bodies that we sampled, these filters rapidly became clogged with sediment. We replaced clogged filters as needed after the filter had processed at least 300 mL of water. This minimum requirement took more processing time but ensured a minimum water sample for each filter. All filters from a sample (range = 2-7) were then processed for qPCR-based eDNA detection for each of the two individual water samples at each site. We only filtered 3000 mL from each sample, which left approximately 800 mL of water at the bottom of the flask where the heaviest sediment load settled. After filtration, filters were folded and stored in 95% ethanol to preserve the samples until DNA extraction. We used the same technique to filter water from a tank of laboratory-cultured NZMS as a positive control to ensure the qPCR protocols were detecting NZMS DNA. As an additional negative control to check for contamination during the filtration process, we used the same approach described above to filter deionized water in the laboratory to ensure sanitization of the filtering equipment was adequate. Any evidence of *P. antipodarum* DNA in this negative control would indicate that contamination had occurred.

### DNA Extraction and qPCR

Although qPCR was used to estimate NZMS population density by Goldberg *et al*. (2013), our goal was different: detect new NZMS invasions. Accordingly, we used qPCR-based detection of eDNA only to determine NZMS presence/absence. We extracted DNA from the filters and processed the filters for quantitative PCR (qPCR) at University of Iowa. Before DNA extraction began, we used DNAZap as well as the 50/50 bleach mixture to clean the bench space. We then used fresh paper towels on the bench and newly bleached equipment for each sample. We extracted DNA from the filters with a DNeasy Blood and Tissue kit with QIAshredder following the DNA extraction protocol described at the Goldberg lab website (https://labs.wsu.edu/goldberglab/edna-assays). To detect contamination during the extraction and qPCR procedures, we processed an unused filter alongside the filters used for water samples, field and filtering negative controls, and positive control.

We used a fume hood that was sterilized with 50/50 bleach and DNAZap for loading qPCR plates. The forward primer sequence was TGTTTCAAGTGTGCTGGTTTAYA, the reverse primer sequence was CAAATGGRGCTAGTTGATTCTTT, and the probe sequence was 6FAM-CCTCGACCAATATGTAAAT. These primers were designed to amplify a polymorphic section of cytochrome *b* (Goldberg *et al*. 2013). The probe was designed to have no ambiguous bases, and the probe sequence was shown to work for NZMS without providing false positives in the presence of the commonly co-occurring pebblesnail (*Fluminicola hindsii*; Goldberg *et al*. 2013). We used 0.4 uM of each primer and 0.2 uM of the probe along with 1X mastermix and 2.5 uL of DNA extract in 20uL reactions in a Roche LightCycler 480. Cycles began at 95°C for 15 minutes followed by 50 cycles of 94°C for 60 seconds and 60°C for 60 seconds. Following Goldberg *et al*. (2013), amplification was considered positive, and therefore indicated detection of NZMS eDNA, if probe fluorescence reached a phase of exponential increase. Wells without exponential increase in fluorescence were considered negative results. Because of varying sediment load across samples, and, accordingly, varying challenges with clogged filters, the number of filters processed per sample were not equal (N = 2-7 filters per sample). To accommodate this variable, we compared site results by calculating the ratio of the number of DNA amplifications (exponential probe fluorescence phase observed during a given qPCR assay) over the total number of qPCR assays for that sample (N = 6-21 qPCR assays per sample). This ratio is hereafter called detectability, a commonly used metric in eDNA-based detection studies (Jerde *et al*. 2011, Dejean *et al*. 2012, Goldberg *et al*. 2013, Deiner and Altermatt 2014).

If any site previously not known to be invaded by NZMS returned a non-zero detectability, we planned to return to the site to perform a thorough search for physical evidence of NZMS presence. This additional line of evidence was crucial for a field demonstration of the viability of eDNA-based early detection of a new NZMS invasive population.

## RESULTS

Detectability of 0.0 for all negative controls (Table 1) shows that none of the five negative control samples tested positive for *P. antipodarum* eDNA. The laboratory positive control taken during this round returned a detectability of 1.0: the one filter from the one sample tested positive for *P. antipodarum* eDNA (3/3 qPCR assays). Sites at Sixmile Run (CE9; Figure 1) and Cedar Run (CR1) both had detectability of 0.0, indicating no NZMS presence. These negative results at CE9 and CR1 were consistent with the failure to detect *P. antipodarum* in earlier physical searches for the snails. One site of Little Juniata River (Hu1) showed a detectability of 0.0, while another (Hu8) had a detectability of 0.167 (6/36 qPCR assays). Juniata River (Bl9) had a detectability of 0.030 (1/33 qPCR assays) and Yellow Creek (YC1) had a detectability of 0.056 (1/18 qPCR assays). One site on Bald Eagle Creek (PA16) had a detectability of 0.100 (3/30 qPCR assays). These sites with relatively low detectability still indicate NZMS DNA was likely present due to the consistent negative control results, but could indicate an unlikely drift event from another site within the watershed or an error not caught by the controls. However, NZMS may be in these streams but remain undetected, and the low detectability is due to their low density levels rather than error. Returning to these sites in the future will be the first step in determining the cause of this low detectability score.

The highest detectability from a site where *P. antipodarum* had not previously been seen was 0.933 (14/15 qPCR assays) from a site on Bald Eagle Creek (PA27) approximately 5.5 km downstream of where Spring Creek empties into Bald Eagle Creek. This result is strongly suggestive of the physical presence of an invasive NZMS population at PA27. While unusually high water levels prevented a follow-up search for NZMS immediately following the results of the May 2018 study, we were able to return to Bald Eagle Creek at the PA27 location in November 2018 and positively identified a single individual NZMS after thorough searching.

## DISCUSSION

We were able to detect eDNA at a site, PA27, at which *P. antipodarum* had never been seen and that was later confirmed to harbor NZMS (Levri *et al*. 2020). We successfully applied eDNA-based methods for early detection of a previously unknown NZMS invasive population. We also demonstrated that eDNA was able to detect NZMS present at likely very low frequency where previous traditional surveys had not identified their presence. We also found evidence that NZMS eDNA may be present at four other sites (Bl9, YC1, PA16, Hu8) at low detectability, indicating NZMS presence but without physical confirmation to date. An obvious next step is to return to these sites where eDNA was detected but snails have not yet been found. Continuing to monitor these locations using eDNA will allow us to track population density increases in known populations and locate dispersal events as they occur.

DNA of other freshwater species has been detected via eDNA-based approaches up to 12 km downstream from its source (Deiner and Altermatt 2014). These reports are difficult to reconcile with more recent studies suggesting that eDNA does not travel further than ∼200m (Wilcox *et al*. 2016, Bedwell and Goldberg 2019). Resolution is hinted at by an experimental study (Shogren *et al*. 2017) demonstrating that eDNA dispersion is both complex and influenced by stream properties like turbulence and substrate structure to a similar or even greater extent than invader population density and DNA release rates. Future attempts to locate and track low-density NZMS populations will need to address how density of nearby populations and properties of the stream (*e*.*g*., morphology, substrate, and flow) may affect detectability of NZMS in the region.

A future study comparing genotypes of this new invasive population at PA27 to other nearby NZMS populations may be able to identify the source populations (Clusa *et al*. 2016) and address the possibility of DNA drift influencing detectability at PA27. In particular, finding that the genotypes of eDNA detected at PA27 match both the Spring Creek population upstream and the individuals found physically at PA27 would mean that we could not formally rule out a scenario where we had detected drifting DNA from a different site. Additional insight into the role of drifting DNA will also come from experiments aimed at characterizing the distance and time over which drifting NZMS DNA specifically can be detected in the environment. This information on drifting DNA will also set the stage for the exciting possibility of identifying the presence of NZMS from sections of the watershed downstream from areas they have invaded. If so, monitoring watersheds by sampling the river they flow into near the mouth and moving upstream may be a way to rapidly survey larger regions for invaders.

While returning to the PA27 site to search for NZMS was our highest priority because of the very high detectability, sites with low detectability still require thorough searches to see if our detection of NZMS eDNA at these locations might also be linked to previously unknown invasions. Given the fact that only one individual was found at PA27, these other sites may harbor NZMS at population densities still too low to find via traditional search methods. We intend to return to these sites for continued traditional searches for physical evidence of the presence of NZMS. We also intend to include these sites in future eDNA surveys, and will focus on later (vs. earlier)-season efforts, which maximizes detectability in river systems (Bedwell and Goldberg 2019). We predict that these sites will either again test positive for *P. antipodarum* (reflecting an established and likely increasing population; in this case, detectability should increase) or will return negative results. The latter would either implicate a rare drifting DNA event and/or suggest that contamination (or mismatches with another species’ DNA; Wilcox *et al*. 2013) might not have been totally eliminated from our procedures.

Our results, along with the recent discovery of NZMS in the Bald Eagle Creek in Lock Haven (Levri *et al*. 2020) indicate that NZMS has expanded its invaded range in central Pennsylvania beyond the Spring Creek watershed. The discovery of NZMS in Spring Creek in 2013 was of great concern due to the potential for spread of the snail from that location to other streams and watersheds across the Appalachian Mountains and the eastern US because of the popularity of Spring Creek for trout fishing. Given the widespread success of NZMS in the western US, it would follow that within a few years there would be additional discoveries of NZMS in the Mid-Atlantic region (Simberloff 2014). Once NZMS establishes in a new location it would likely take a few years for the population to become large enough to be detected via traditional approaches (Simberloff 2014, U.S. Department of the Interior 2016). Now, a few years after its initial discovery at Spring Creek in Pennsylvania, there has been an increase in new findings of the snail in the eastern US. In 2017, NZMS was found in two streams in Syracuse, NY and in the Gunpowder Falls River in Maryland. In 2018, NZMS was discovered in the Musconetcong River in New Jersey and Little Lehigh Creek near Allentown, PA (R. Morgan, pers. comm.), and in 2020, it was found in Codorus Creek near York, PA (C. Urban, pers. comm.). Unfortunately, these populations are all very well established and may already have sourced new unknown invasions elsewhere. That newly invaded sites can now be detected before NZMS populations are large will allow more rapid measures to be taken to educate the public and limit their spread.

In particular, we can use eDNA approaches to slow the rate of NZMS spread in the eastern US by making recreational water users aware of NZMS presence, implementing checkpoint procedures near invaded sites, and ideally, limiting access to invaded locations as a rapid response (Simberloff 2014, U.S. Department of the Interior 2016). We have provided a proof-of-principle for using eDNA to detect previously unknown populations of NZMS. Future studies should focus on smaller water samples for more rapid filtration and sample more frequently and broadly across the region in order to detect establishing populations and monitor established invasive population densities (Goldberg *et al*. 2013). There is also the exciting opportunity for citizen science to contribute to these efforts (Biggs *et al*. 2015). With the cooperation of those individuals that use these ecosystems for recreation, it may be beneficial to pursue a program wherein sterile containers and instructions are supplied to fishermen, boaters, and kayakers. After sampling, the containers can be dropped off at a laboratory for processing. While such a program could introduce more chances for error, continual assessment of potential invasion sites would allow for cross-referencing results. This distributed method of early detection would cast a wide net in which to catch these destructive aquatic invaders.

## DECLARATIONS

### Author’s contributions

James Woodell planned the project, adapted eDNA techniques for the project, led sampling and analysis, and was the primary author of this manuscript. Dr. Neiman helped plan the project and helped write the project proposal and final manuscript. Dr. Levri led the project proposal, provided assistance with sampling, and oversaw the project and final manuscript.

All authors have approved this manuscript for pre-print release and it has not yet been accepted for publication elsewhere.

## Acknowledgements

Mid-Atlantic Panel on Aquatic Invasive Species for project funding. Dr. Caren Goldberg for answering our sampling and sample processing questions. Dr. Gery Hehman for his training and assistance with the use of equipment at the University of Iowa’s Carver Center for Genomics. Elaine Vizka for her GIS expertise.

